# Performance of Chrysanthemum Cultivars under Agro-climatic condition of Chhattisgarh plains

**DOI:** 10.1101/2023.11.28.568964

**Authors:** Madhusmita Bhoi, Tarsiyus Tirkey

**Author notes:** Email: [, ].

## Abstract

Chrysanthemum is the second largest flower after rose in the global ornamental plants market; It is third in terms of cut flower and ranks fifth in global floriculture trading. Therefore, floricultural study of chrysanthemum has gained widespread attention recently. As the state of Chhattisgarh is rising as a floral hub, we are propelled to determine suitable cultivars of chrysanthemum for different characters in Chhattisgarh plains. To this end, we conducted an experiment based on Randomized Block Design (under field condition) during the year 2016-2017, with seven cultivars of Chrysanthemum namely Pusa Sona, Pusa Arunodaya, Pusa Kesari, Pusa Chitraksha, Pusa Aditya, Mother Teresa, and White Prolific. Our results indicate that Pusa Sona exhibits the early bud appearance (32.80 days), early flowering (60.27 days), maximum number of flowers per plant (137.80), and maximum number of branches per plant (19.73). Further, maximum plant height (49.47 cm) and plant spread (32.63 cm) were observed in Pusa Chitraksha. While Pusa Kesari recorded maximum flower diameter (7.00 cm) and maximum weight of flowers per plant (177.50 g), the maximum flowering duration (59.40 days) was noted under Pusa Aditya. We are certain that these novel findings would help floriculturists’ decision making significantly regarding chrysanthemum plantation.

## INTRODUCTION

With the advancement of technologies, floriculture yields have grown manifolds. This has made floriculture a lucrative entity for countries worldwide. Particularly, Chrysanthemum is a frontrunner for its diversity in terms of color, shape, and sizes. The genus Chrysanthemum belongs to Asteraceae family comprises of huge biodiversity in their growth habitat, flowering behavior, flower and foliage color, shape and size. This genus includes over 200 species of annuals and herbaceous perennials. The word “Chrysanthemum” comes from two Greek words, Chrysos – golden and Anthos - flower which means golden flower. They were named so because of the golden color of its original flower. It is the second largest flower after rose among the ornamental plants traded in the global flower market (Singh & Karki). In terms of cut flower, Chrysanthemum ranks third, and as a pot plant, it ranks fifth in global trade of floriculture commodities (Singh and Karki). It is widely used as loose flower, cut flower as well as pot plant (Kher, 1988). In addition, it is a major flower for making garlands, veni, bouquets and for worship. In the last two decades, chrysanthemum has become a leader of commercial flower crops of several countries. It Unsurprisingly, in India as well, Chrysanthemum is a flag holder both as a commercial crop and in flower exhibitions. However, there are inherent challenges to further support the rising demand of Chrysanthemum as discussed next.

The performance of Chrysanthemum cultivars varies with region, season and other growing conditions (Tomar *et al*. 1972). As a result, cultivars, which perform well in one region, may not perform well in other regions of varying climatic conditions (Gupta and Datta, 2005). The quest for selecting suitable high yielding variety for a Tomer region leads to the requirement of collection and evaluation of available cultivars. Moreover susceptibility of existing cultivars to different biotic stresses and condition augments the need of promising cultivars. The ultimate yield and production of quality flower and resistance to biotic factors depend upon the selection of suitable cultivars for a particular locality. Therefore, it is necessary to identify the most suitable cultivars for a certain agroclimatic condition which may beneficial for flower growers. In this context, we devise experiments to identify the appropriate cultivars of Chrysanthemum for our state, i.e., Chhattisgarh.

## MATERIALS AND METHODS

We choose seven cultivars of chrysanthemum namely Pusa Sona, Pusa Arunodaya, Pusa Kesari, Pusa Chitraksha, Pusa Aditya, Mother Teresa, and White Prolific for our study. A field experiment was carried out to evaluate their performance at the Horticultural Research cum Instruction Farm of the Department of Floriculture and Landscape Architecture, College of Agriculture, Indira Gandhi Krishi Vishwavidyalaya, Raipur, Chhattisgarh, during the year 2016-2017. Raipur is situated in the central part of Chhattisgarh at a latitude of 21^0^16^’^ N and longitude of 81^0^36^’^ E, and at an altitude of 289.56 meters above the mean sea level. We perform the experiment in Randomized Block Design (RBD) with 3 replications. Five plants were randomly selected from each treatment for carrying out evaluation studies. All the recommended cultural practices were followed. The data on various vegetative characters and floral characters were, recorded and statistically analyzed.

## RESULTS AND DISCUSSION

### Vegetative characters

Cultivar significantly differed for plant height, branches and plant spread (Table 1). The study revealed that maximum plant height was recorded in Pusa chitrakhsha (49.47 cm) whereas minimum plant height was recorded in Pusa Sona (19.19 cm) variety. Pusa Sona recorded maximum number of branches per plant (19.73 cm) and minimum number of branches per plant was obtained in Pusa kesari (8.60 cm). Maximum plant spread was found in Pusa Chitraksha (32.63 cm) Whereas, minimum plant spread was observed in White prolific (19.32cm). Differences in vegetative characters can be attributed to several factors including genetic and climatic like soil condition, temperature, light, nutrition, etc. During the experiment since all the cultivars were treated with similar climatic condition, the differences in plant height can be attributed mainly due to variation in genetic character (Shirohi and Behra, 2000). Similar findings have been reported by Vetrivel *et al*. (2014), Srilatha *et al*. (2015).

**Table 1:** Evaluation of Chrysanthemum varieties for vegetative characters:

| Varieties | Plant height (cm) | No. of branches per plant | Plant spread (cm) |
| --- | --- | --- | --- |
| T<sub>1</sub>=Pusa Sona | 19.19 | 19.73 | 19.75 |
| T<sub>2</sub>=Pusa Arunodaya | 29.85 | 10.53 | 22.16 |
| T<sub>3</sub>=Pusa Kesari | 39.88 | 8.60 | 28.73 |
| T<sub>4</sub>=Pusa Chitraksha | 49.47 | 12.13 | 32.63 |
| T<sub>5</sub>=Pusa Aditya | 36.57 | 15.47 | 28.86 |
| T<sub>6</sub>=Mother Teresa | 34.91 | 11.13 | 30.11 |
| T<sub>7</sub>=White Prolific | 39.31 | 13.33 | 19.32 |
| SEm± | 0.82 | 0.64 | 0.63 |
| CD at 5% level of significance | 2.56 | 1.98 | 1.95 |

### Flowering Attributes

From the Table 2, it was apparent that, significantly early flower bud appearance (32.80 days) and early flowering were recorded in Pusa Sona (60.27 days). While, the maximum number of days for first flower bud appearance (50.80 days) and first flower opening (80.13 days) were recorded in Mother Teresa. The variation for early or late bloom seems to be the varietal character which is in agreement with the findings of Kanamadi and Patil (1993), Behera *et al*. (2002) in chrysanthemum.

**Table 2:** Evaluation of Chrysanthemum varieties for flowering attributes:

| Varieties | Days to bud initiation | Days to flowering | Flower Diameter (cm) | Number of flowers per plant | Weight of flowers per plant (g) | Flowering Duration |
| --- | --- | --- | --- | --- | --- | --- |
| T<sub>1</sub>=Pusa Sona | 32.80 | 60.27 | 3.11 | 137.80 | 67.05 | 57.73 |
| T<sub>2</sub>=Pusa Arunodaya | 35.07 | 66.00 | 6.12 | 37.40 | 102.38 | 48.60 |
| T<sub>3</sub>=Pusa Kesari | 40.67 | 69.07 | 7.00 | 42.40 | 177.50 | 50.00 |
| T<sub>4</sub>=Pusa Chitraksha | 43.93 | 75.67 | 5.87 | 100.27 | 136.59 | 56.93 |
| T<sub>5</sub>=Pusa Aditya | 35.53 | 62.73 | 5.04 | 85.67 | 94.48 | 59.40 |
| T<sub>6</sub>=Mother Teresa | 50.80 | 80.13 | 3.90 | 134.53 | 82.05 | 46.47 |
| T<sub>7</sub>=White Prolific | 41.93 | 74.00 | 5.64 | 42.53 | 140.36 | 42.13 |
| SEm± | 1.04 | 1.37 | 0.24 | 1.46 | 1.62 | 0.86 |
| CD at 5% level of significance | 3.24 | 4.27 | 0.75 | 4.55 | 5.04 | 2.67 |

Pusa Kesari recorded maximum flower diameter (7.00 cm) Whereas, the minimum flower diameter was observed in Pusa Sona (3.11cm). Maximum number of flowers per plant was found in Pusa Sona (137.80) which was at par with Mother Teresa (134.53) and minimum number of flowers per plant was found in Pusa Arunodaya (37.40). Maximum weight of flowers per plant was observed in Pusa Kesari (177.50 g) and minimum weight of flowers per plant was found in Pusa Sona (67.05 g). Pusa Aditya recorded maximum flowering duration (59.40 days) which was at par with Pusa Sona (57.73 days), Pusa Chitrakhsha (56.93 days) Whereas, the minimum flowering duration was obtained in White Prolific (42.13 days).Significant differences were recorded in all the varieties for flowering attributes which is in agreement with the findings of Deka and Paswan (2001), Jayanthi and Vasanthachari (2003), Balaji *et al*. (2004) and *Dilta* et al. (2005) and Manohar Rao (2003) and Pratap (2006) in chrysanthemum.

## Conclusion

Chrysanthemum has become a leader of commercial flower crops of several countries including India, recently. However, the various cultivars of chrysanthemum vary region-wise in terms of production shape, sizes, and color. To this end, we perform experiments with its seven cultivars namely Pusa Sona, Pusa Arunodaya, Pusa Kesari, Pusa Chitraksha, Pusa Aditya, Mother Teresa, and White Prolific for the region of Chhattisgarh. Our randomized block design-based experiments show that the variety Pusa Sona was found early in the Chhattisgarh agro climatic condition followed by Pusa Aditya which can help to fetch the early market. Pusa Sona also recorded maximum number of flowers per plant, and also maximum number of branches per plant. Pusa Aditya recorded maximum flowering duration followed by Pusa Sona. Due to its specification of maximum flowering duration, these flowers can be attributed for longer period of availability in the market. From this study, we concluded that Pusa Sona is the most promising cultivar in Chhattisgarh agroclimatic condition followed by Pusa Aditya. We plan to carry out similar experiments in other states of India such as Odisha, West Bengal, And Bihar to devise a holistic region-wise scheme for chrysanthemum crop.

## Notes

### Competing Interest Statement

The authors have declared no competing interest.

## Reference

Deka, K. K., Paswan, L. 2001. Growth performance of some standard chrysanthemum (Dendranthema grandiflora Tzelev) cultivars under the agro climatic condition of Jorhat. Research-on-crop.,2 (3): 364–367.

Joshi, M., Verma, L. R., and Masu, M. M., .2009. Performance of different varieties of chrysanthemum in respect of growth, flowering and flower yield under north Gujarat condition The Asian Journal of Horticulture., 4 (2): 292–294.

Kanamadi, V. C and Patil, A. A 1993. performance of chrysanthemum varieties in the transitional tracts of Karnataka. South Indian Hort., (41): 58–60.

Rao, Puneth, Parul, V. K. and Sharma, S. K. 2011. Evaluation of different chrysanthemum (Chrysanthemum moriflolium) genotypes under mid hill condition of Garhwal Himalaya. Indian Journal of Agricultural Sciences, 81(9): 830–833.

Rathore, I., Kaushik, R. A., Upadhyay, B. and Mahawer, L.N.2016 varietal evaluation of chrysanthemum (Chrysanthemum morifolium ramat.) Progressive Research – An International Journal., 11 (Special-I): 283–289

Srilatha, V., Kumar, K. S., and Kiran,Y. D. 2015 Evaluation of chrysanthemum (Dendranthema grandiflora T.) varieties in southern zone of Andhra Pradesh. Agric. Sci. Digest., 35 (2): 155–157.

Suvija, N.V., Suresh, J., Kumar, S. R. and Kannan, M.2016 Evaluation of Chrysanthemum (Chrysanthemum Morifolium Ramat) Genotypes for Loose Flower, Cut Flower and Pot Mums International Journal of Innovative Research and Advanced Studies (IJIRAS).,3(4): 2394–4404

Vetrivel,T. and Jawaharlal, M. 2014. Evaluation of Chrysanthemum (Dendranthema grandiflora T.) Varieties for Yield and Quality under Subtropical Hills. Trends in Biosciences., 7(14): 1812–1815.

